## Supplementary figures and images for "Cellular characterization of the mouse collecting lymphatic vessels reveals that lymphatic muscle cells are the innate pacemaker cells"

### Supp Figure 1

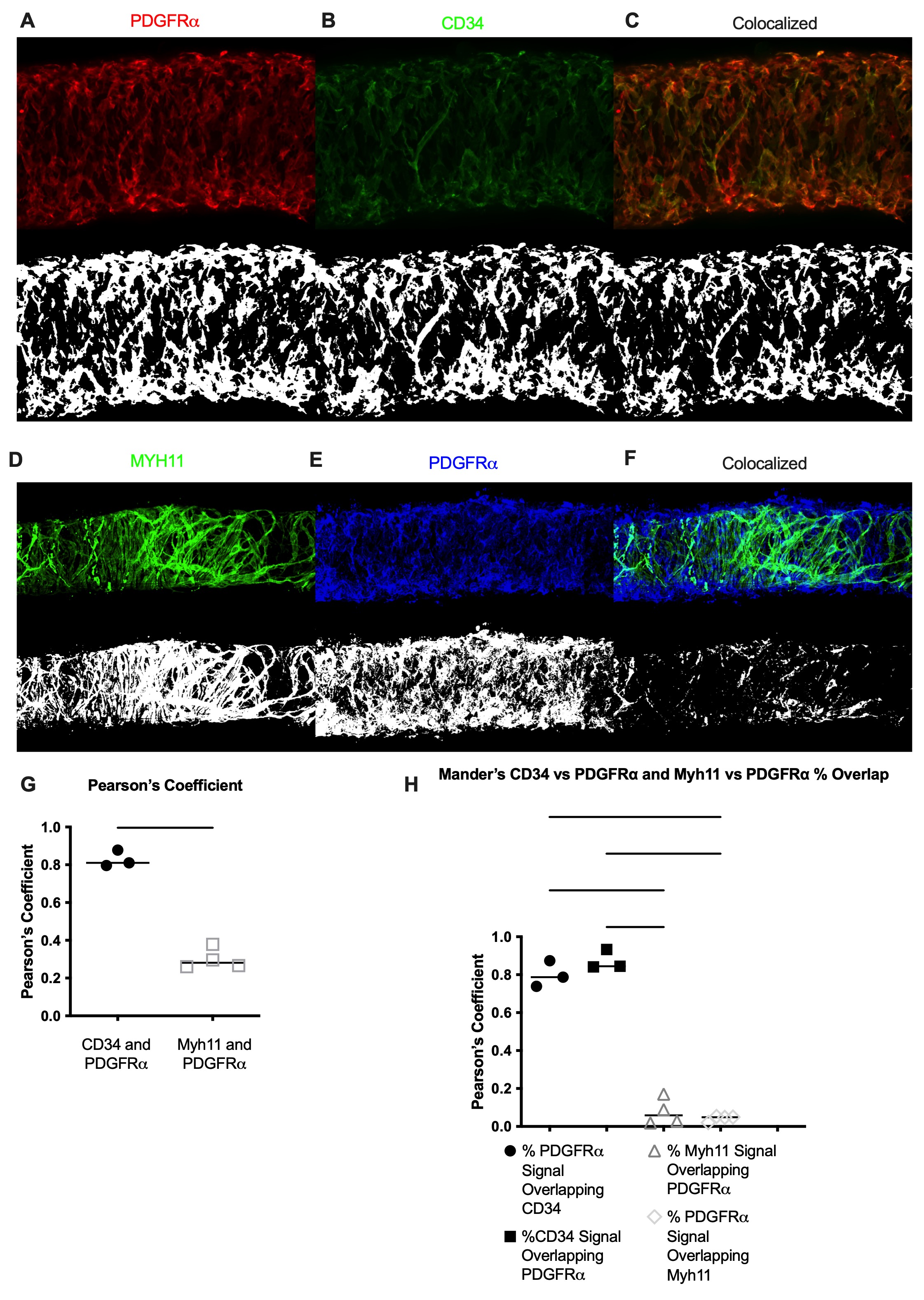

### Supp Figure 2

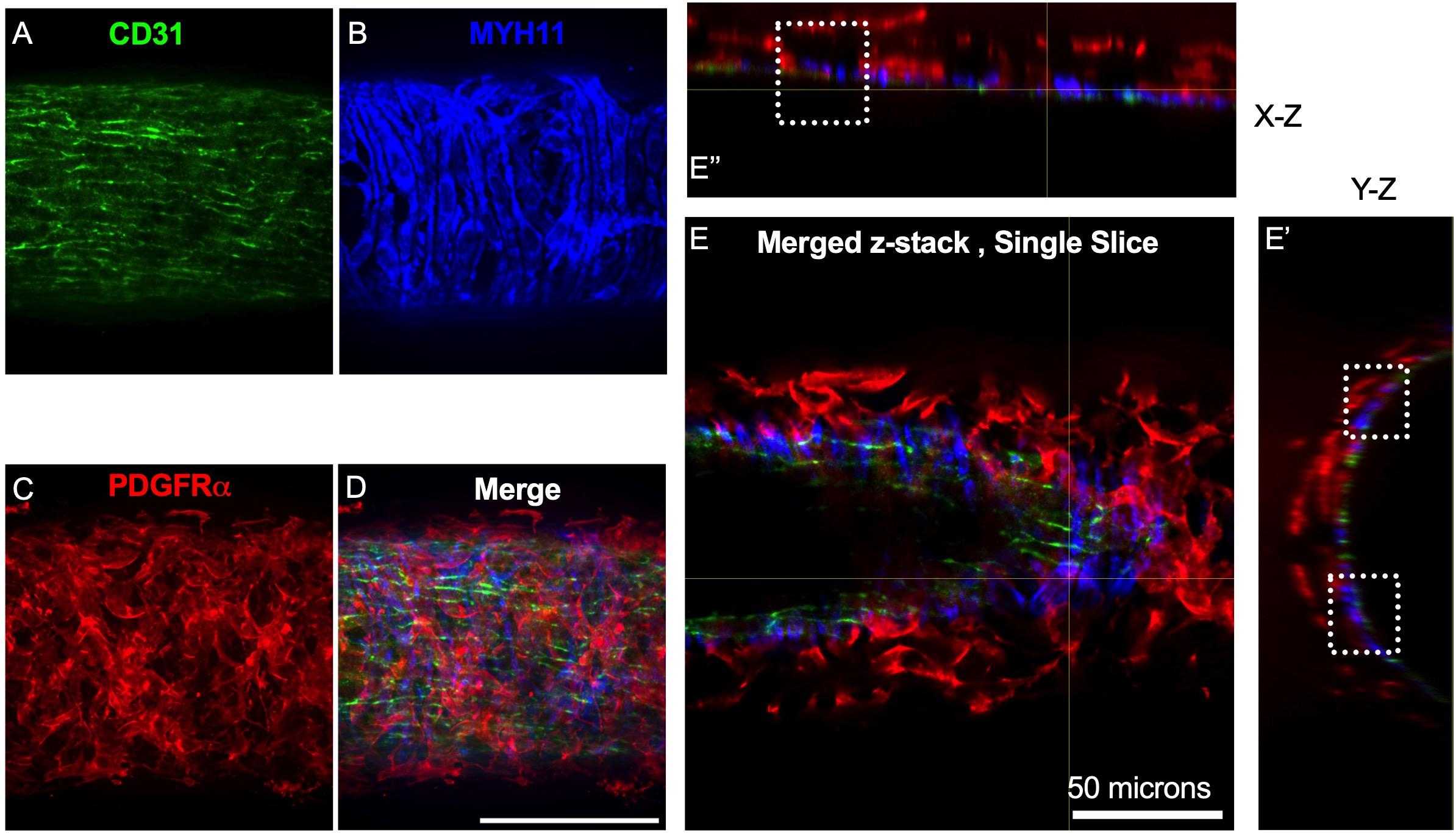

### Supp Figure 3

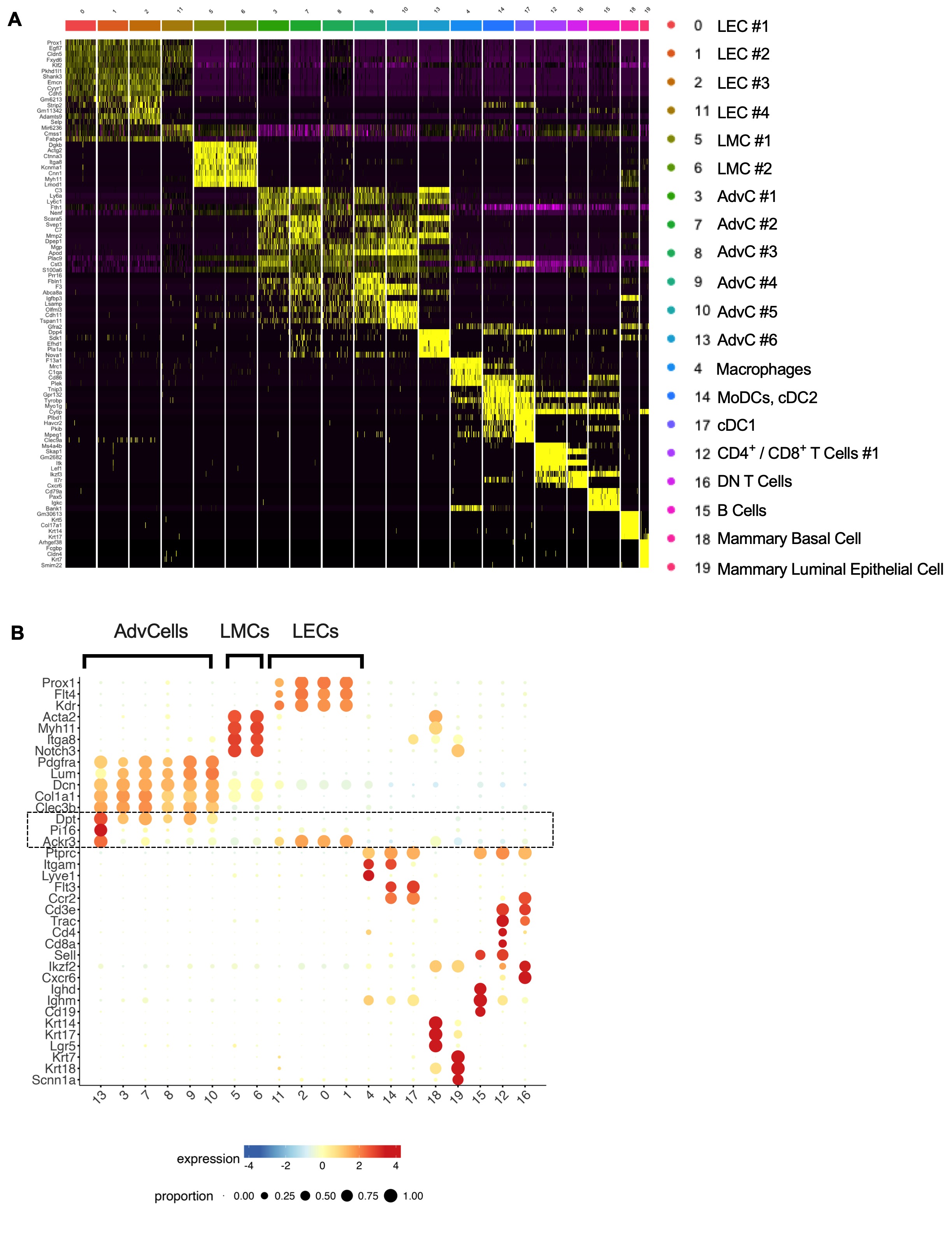

### Supp Figure 4

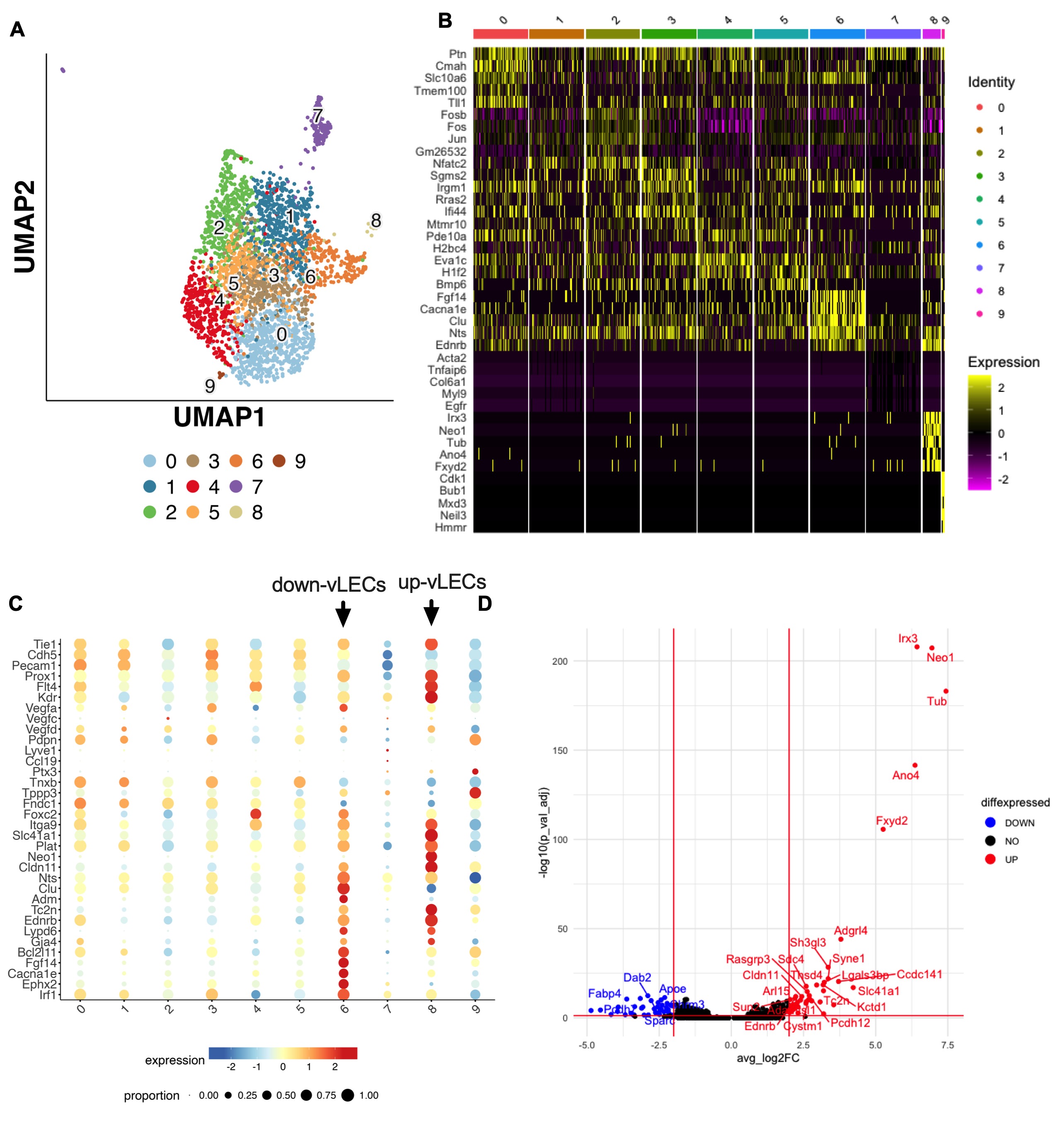

### Supp Figure 5

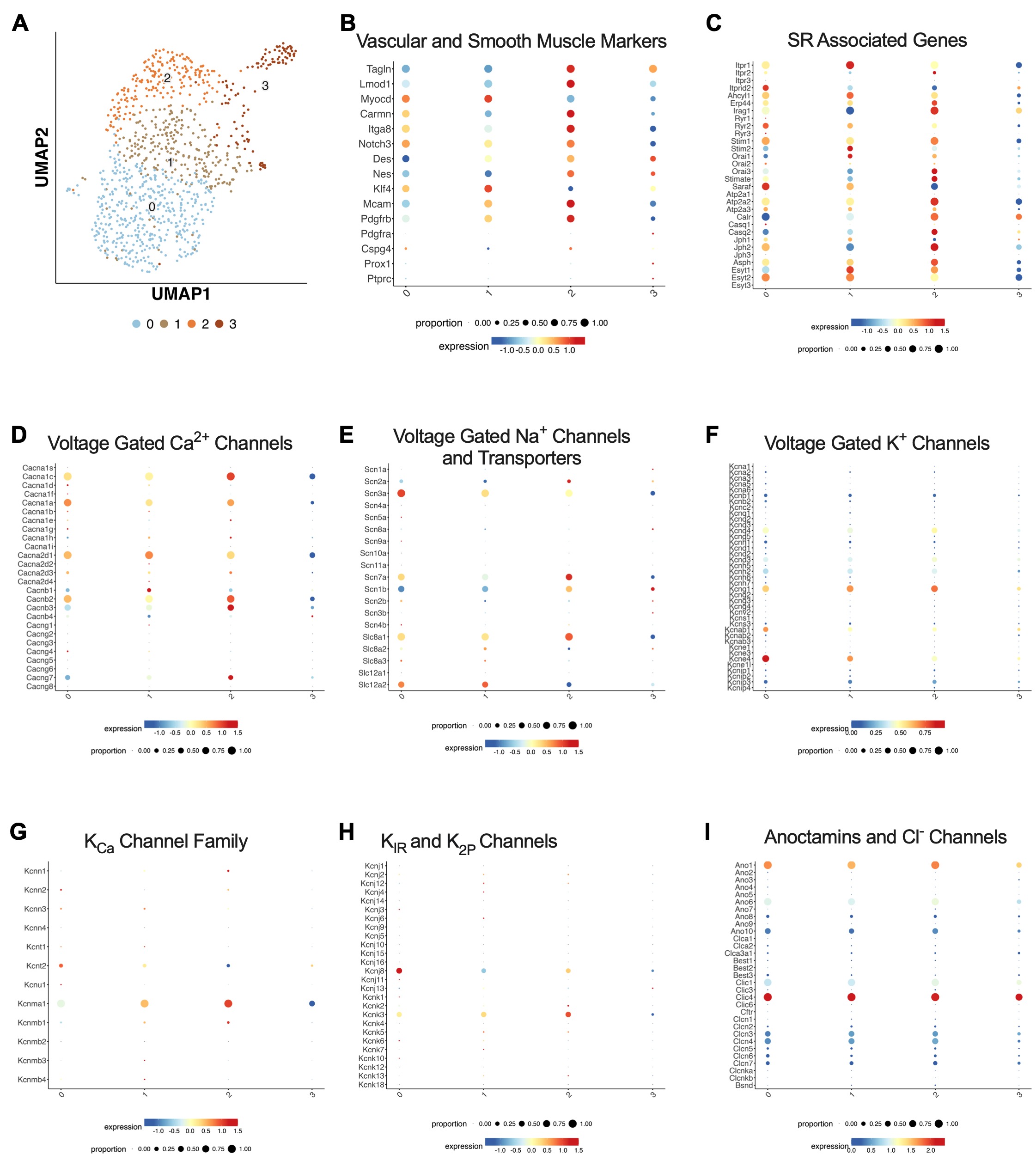

### Supp Figure 6

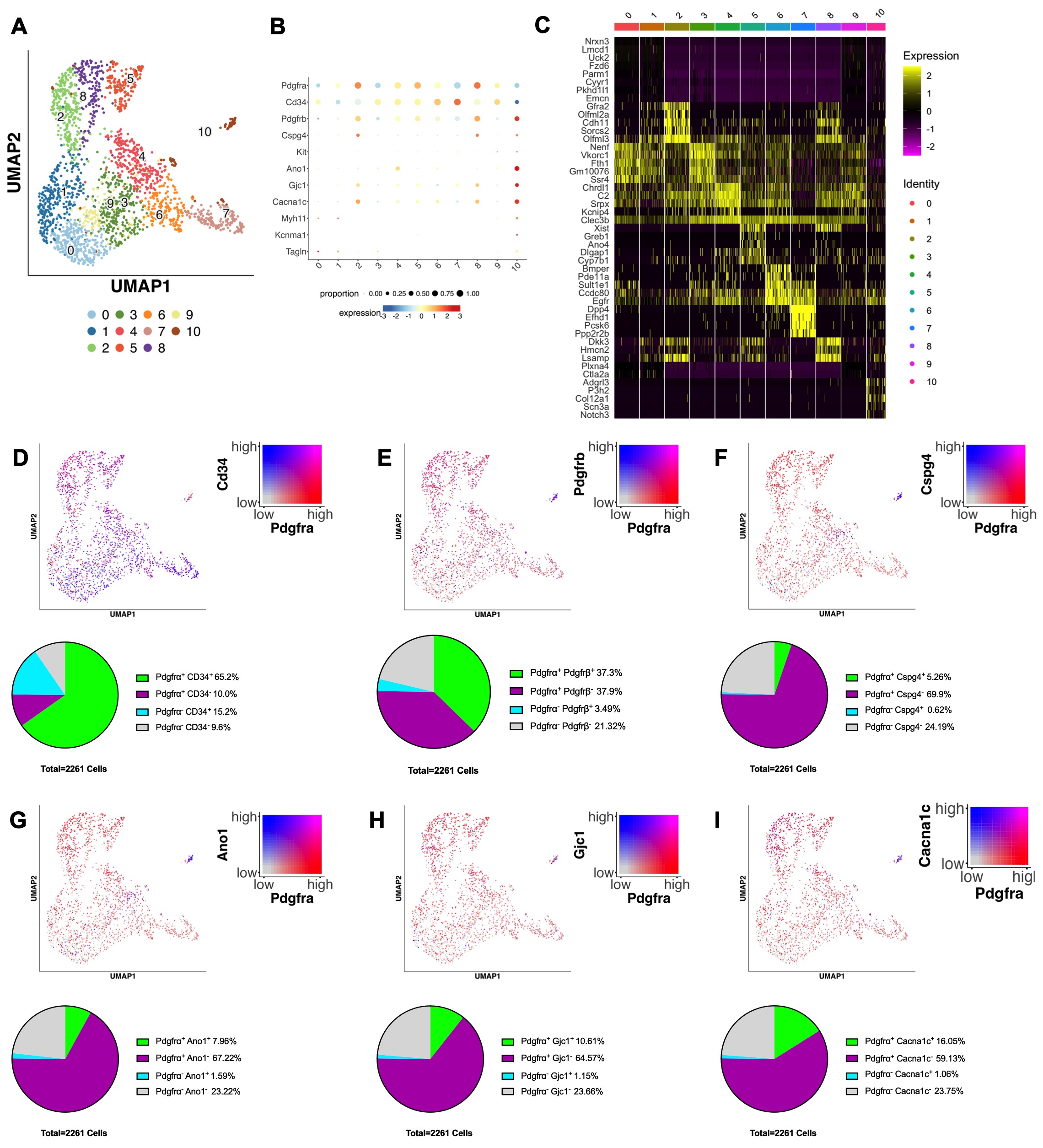

### Supp Figure 7

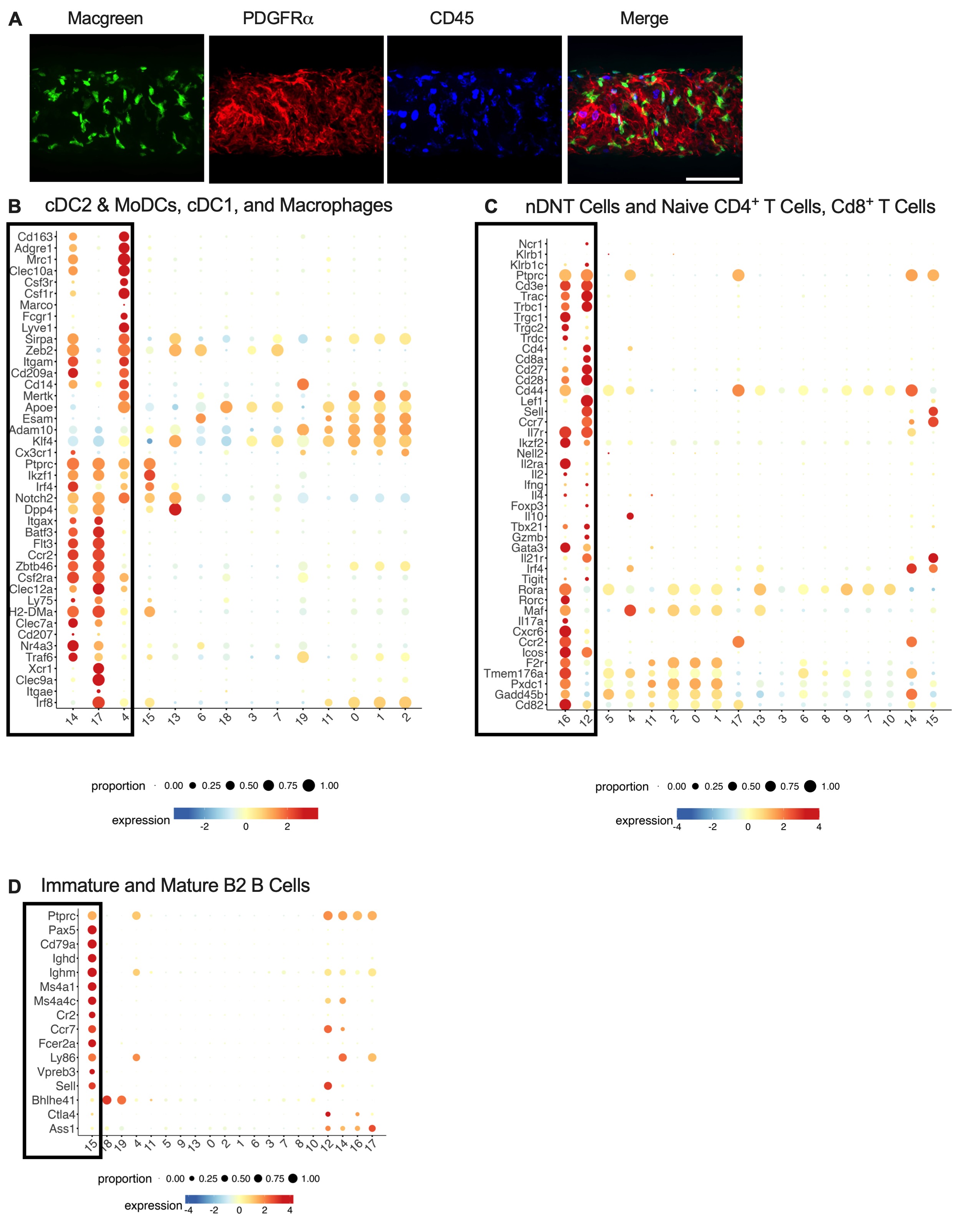

### Supp Figure 8

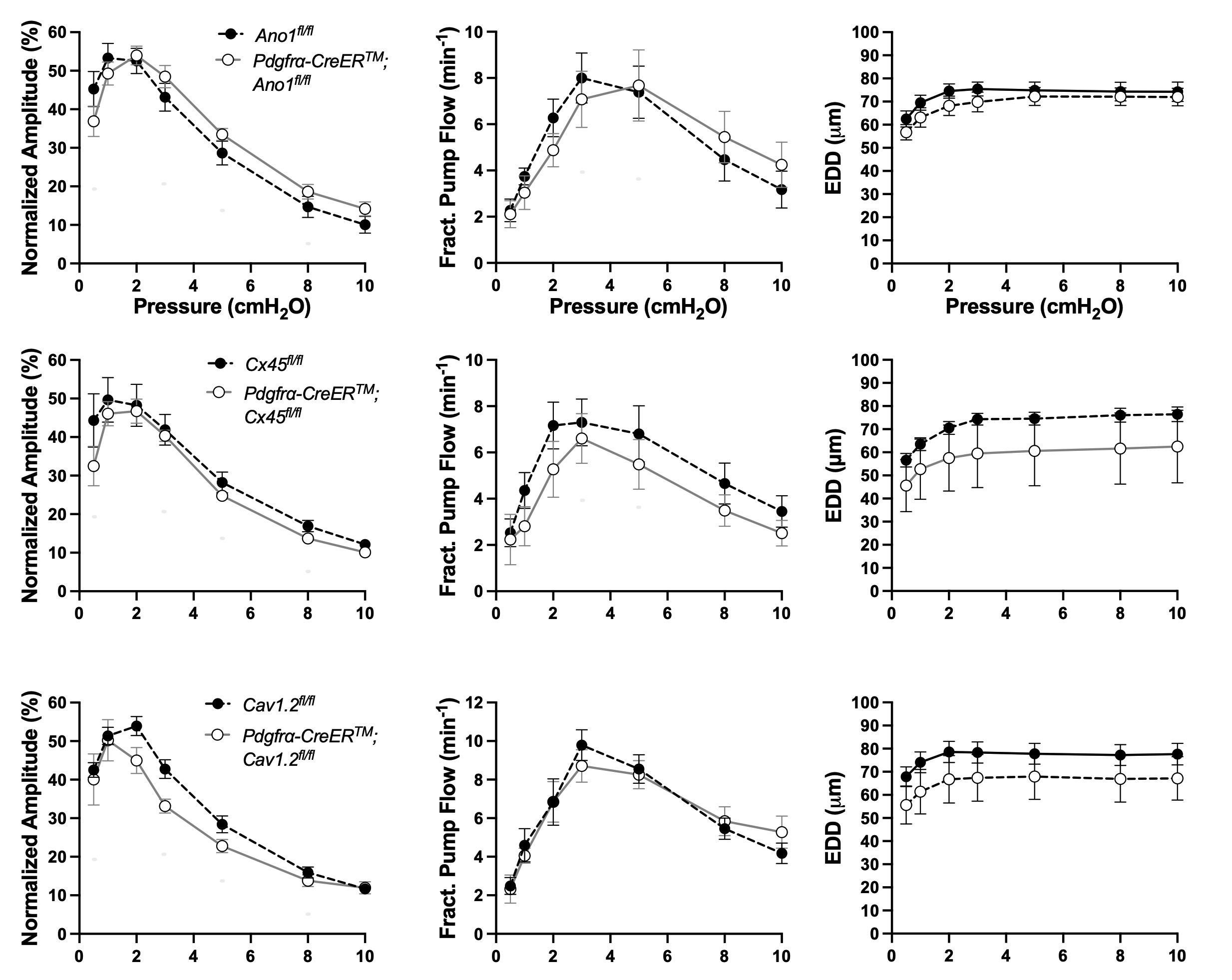

### Supp Figure 9

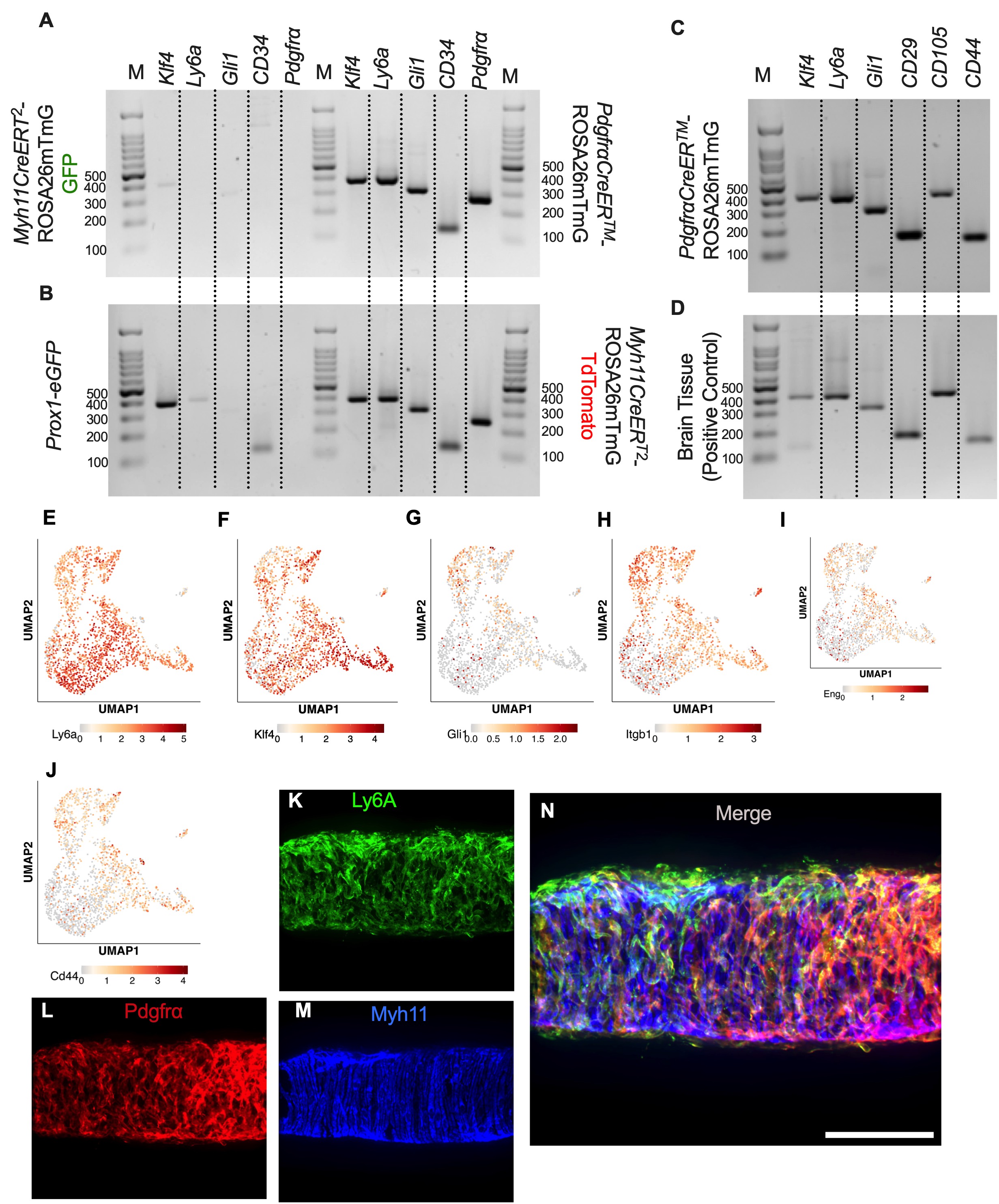
